# Capturing the missing gene content of elite wheat by analysis of wheat landraces

**DOI:** 10.64898/2026.07.30.741942

**Authors:** Teng Li, Jack Bruton, Felipe E. Albornoz, Mitchell Bestry, Aria Dolatabadian, Tessa MacNish, Thomas Bergmann, Philipp Bayer, Jacqueline Batley, David Edwards

## Abstract

There is a need to breed superior wheat cultivars to meet increasing global food demand. However, modern cultivars have undergone a substantial loss of genetic diversity due to intensive breeding. A pool of genetic diversity remains untapped in wheat landraces and analysis against a single reference cannot detect genes that the reference lacks. Pangenomics can help identify genes of potential agronomic importance in these old lines that can be applied to accelerate wheat improvement. Here, we have constructed the largest wheat pangenome to date, representing 1,061 diverse individuals (827 landraces and 234 modern cultivars) from 47 countries across six continents. We identified 10,426 predicted gene models specific to landraces that are enriched for functions associated with disease resistance, abiotic stress adaptation, and symbiosis, as well as 176 landrace-specific candidate disease resistance loci. Phylogenetic analysis inferred from gene presence/absence variation reflects SNP-based ancestry-associated structure, while independent Iso-Seq data from 14 cultivars showed 96.4% concordance with cultivar-specific gene presence calls from our PAV analysis, supporting the robustness of the resource. Our results reveal that landraces harbour an extensive repertoire of genes that are absent from modern wheat cultivars and provide a foundational resource that links this gene content variation to specific accessions and ancestral groups, enabling candidate genes to be prioritised in donor lines for subsequent introgression and supporting the systematic reintroduction of adaptive variation to enhance wheat resilience and sustainability in the face of climate change.

## Introduction

Bread wheat (*Triticum aestivum* L., AABBDD) is one of the most widely cultivated cereal crops worldwide (Liu et al., 2022), supplying approximately one fifth of human caloric intake (Lobell et al., 2011). As global food demand increases, there is a need to enhance crop yield by breeding superior cultivars (Anderson et al., 2020). Comparative genomic studies highlight the large variation in gene content among individuals of the same plant species (Danilevicz et al., 2020; Golicz et al., 2016). This raises the possibility of identifying novel genes for important agronomic traits through sequencing diverse germplasm. Wheat has experienced a substantial narrowing of its ancestral gene pool during domestication, limiting the availability of adaptive genetic variation that could enhance resilience and yield (Blower et al., 2025; Haudry et al., 2007; Montenegro et al., 2017). However, such lost genetic diversity can be reintroduced into modern varieties, as they persist in traditional landraces and wild relatives (Reif et al., 2005; Smale et al., 2002).

Landraces are locally adapted populations maintained by traditional farming that, as a group, harbour high genetic diversity, because each landrace reflects a unique evolutionary history shaped by local environmental selection pressures (Dwivedi et al., 2016). Genes from landraces, as well as those from wild relatives, can be used to improve modern crop cultivars by introgressing disease-resistance and stress-tolerance genes (Singh et al., 2018; Toit, 1989; Valdez et al., 2012). For example, McFadden (1930) introgressed the stem-resistance gene *Sr2* into the wheat cultivar ‘Marquis’ using the landrace ‘Yaroslav emmer’ as the donor, producing the modern cultivar ‘Hope’ (McFadden, 1930). Another example is the wheat–rye 1RS.1BL translocation, in which the short arm of rye chromosome 1R was introgressed into wheat chromosome 1B and has a significant impact on wheat yield (Rabinovich, 1998). Similarly, Placido et al. (2013) introgressed a chromosome segment (7DL) from a wild wheat relative into a modern cultivar that improved water stress adaptation (Placido et al., 2013).

Several other studies have introgressed known genes to improve modern wheat cultivars, such as the *1Ay21\** allele that increases protein content and yield (Roy et al., 2020), and two genes (*Sr22* + *Sr25*), which in combination provide stem rust resistance (Sibikeev et al., 2021). Similar findings have been reported in other crop species. For example, Neeraja et al. (2007) used marker-assisted backcrossing to introgress the submergence tolerance gene *Sub1* from the rice landrace ‘FR13A’ into the modern cultivar ‘Swarna’ (Neeraja et al., 2007). Similarly, Qiao et al. (2016) crossed wild rice with a modern cultivar and identified 25 quantitative trait loci that enhanced multiple agronomic traits (Qiao et al., 2016). These studies highlight the importance of large-scale genetic profiles of landraces and wild relatives for capturing global genetic diversity and identifying genes of agronomic importance (Barbosa et al., 2021; Hao et al., 2020; Zhang et al., 2022).

Pangenomes have substantially advanced our understanding of genome-wide variation (Bayer et al., 2020; Hu et al., 2025) and its role in crop domestication and breeding (Li et al., 2025; MacNish et al., 2026; Schreiber et al., 2024). For example, a rice pangenome assembled from 129 wild and 16 cultivated genomes identified 13,728 genes exclusive to wild rice, revealing a greater abundance and diversity of resistance-gene analogues (RGAs) in wild relatives than modern cultivars (Guo et al., 2025). Recent pangenome studies in *Brassica* species further demonstrate the power of multi-genome resources for identifying novel functional genes and domestication-related variants (Amas et al., 2023; Guo et al., 2024; Ma et al., 2026).

The first wheat pangenome was constructed by Montenegro et al. (2017) using an iterative assembly approach, reassembling the Chinese Spring reference genome alongside whole-genome resequencing data from 18 wheat cultivars (Montenegro *et al*., 2017). This work demonstrated that Chinese Spring is an outlier with substantial sequence differences from other modern cultivars. The subsequent “10+ wheat genomes” project generated ten chromosome-scale and five scaffold-level assemblies from global wheat breeding programs, revealing extensive structural rearrangements, introgressions from wild relatives, and a repertoire of nucleotide-binding leucine-rich repeat (NLR) proteins (Walkowiak et al., 2020). Later, Bayer et al. (2022) built the first wheat graph pangenome and associated visualisation tool, Wheat Panache (Bayer et al., 2022), which enabled accurate identification of the physical locations of genomic and structural variants. More recently, a wheat pangenome constructed using 17 newly assembled chromosome-level genomes from representative Chinese cultivars plus four previously published assemblies, identified 170,517 gene families (Jiao et al., 2025). Although foundational, these studies are limited by relatively small sample sizes, narrow geographic representation, or are focused on elite or region-specific cultivars. As a result, global landrace genetic diversity remains underrepresented in pangenome studies. The Watkins collection hosts 832 bread wheat landrace cultivars from 32 countries acquired by A.E. Watkins in the 1930s (Wingen et al., 2014). These lines capture extensive historical diversity predating modern breeding (Winfield et al., 2018). Recently, whole-genome re-sequencing has been performed on this collection in addition to a selection of modern cultivars (Cheng et al., 2024), which assigned the collection into ancestral groups (AGs) based on phylogeny and population structure using SNP data. Modern wheat cultivars mostly overlap with AG2 and AG5, leaving the remaining five AGs as an extensive reservoir of untapped diversity. A recent genome-wide association study (GWAS) further demonstrated the breeding value of this diversity, identifying extensive variation in stripe rust resistance and 87 quantitative trait loci (QTL) across 297 Watkins accessions, including several potentially novel resistance loci (Singh et al., 2026). The Watkins collection, therefore, offers a unique opportunity for characterising the global genetic diversity of wheat and builds upon previous pangenomes.

Here, we present a bread wheat pangenome to identify genes lost during modern breeding. The pangenome was constructed from 1,061 globally diverse individuals using publicly available data, integrating 21 high-quality genome assemblies with whole-genome sequencing data from 827 Watkins landraces and 217 modern cultivars (four of the resequenced modern cultivars were also represented in the 21 assemblies set). Unlike previous pangenomes that are limited in both diversity and landrace representation, this study brings together broad geographic coverage and deep historical diversity, making it the largest and most geographically comprehensive wheat pangenome assembled to date. Its major value lies in enabling the discovery of genes present in landraces but absent in modern cultivars, highlighting a reservoir of untapped genetic diversity that has been largely overlooked in recent crop improvement. We first constructed a graph-based reference pangenome from 21 high-quality genomes, then iteratively expanded it by incorporating sequences from additional cultivars. This allowed us to capture rare and geographically restricted genetic variation at an unprecedented scale. We further performed gene presence-absence (PAV) analyses, gene ontology (GO) analyses, and evaluated PAV of agriculturally relevant genes across the full set of 1,061 genomes. These analyses revealed a large repertoire of genes present in wheat landraces but absent from current elite cultivars, many of which are potentially involved in agronomically important functions related to biotic stress responses and environmental adaptation. Functional annotation of these genes suggests their potential for wheat improvement, and together with the landraces that host them, provide the basis for empirical assessment of their potential to improve bread wheat while increasing the genetic diversity of modern breeding pools.

## Results and Discussion

### Constructing a wheat graph pangenome

The Watkins wheat pangenome was constructed by first generating an initial graph pangenome from 21 high-quality genome assemblies (Supplementary Table 1). The graph consisted of 4,762,231 segments, with a mean length of 3,452 bp and a maximum length of 315,658,419 bp (Figure S1A). The graph contained 6,894,289 unique edges, with an edge-to-node ratio of 1.45 and a total node sequence length of 16.44 Gb, of which 1.89 Gb (11.51%) was contributed by non-reference nodes, most likely due to substantial wheat genomic content absent from cv. Chinese Spring, as previously observed (Montenegro *et al*., 2017). The transposable elements (TEs) made up 84.43% of the graph genome, of which 65.64% were long terminal repeat (LTR) and 18.04% were DNA transposons (Supplementary Table 2). Across the 21 genome assemblies, the median assembly-to-graph mapping coverage was 94.28% (75.531–99.997%) (Figure S1B). There were 548,654 segments (11.3%) present in all 21 individuals, whereas 524,424 segments (10.8%) were unique to one individual (Figure S1C). In total, 118,038 predicted gene models could be placed in the graph, with Benchmarking Universal Single-Copy Ortholog (BUSCO) analysis indicating a high level of completeness (99.2%). By comparison, the IWGSC RefSeq v1.1 annotation of Chinese Spring contains 107,891 high-confidence gene models (International Wheat Genome Sequencing Consortium (IWGSC), 2018), indicating the additional gene content captured by the graph relative to a single reference genome. Among the 21 individuals, 83,177 gene models (70.5%) were present in all individuals, while 2,488 (2.1%) were unique to one individual (Figure S1D). As the number of genomes increased, the number of core gene models declined, and the pangenome curve approached a plateau when the number of accessions exceeded 15, after which only marginal increases in gene model number were observed (Figure S1E).

### Extending the wheat pangenome with the Watkins landraces

Whole-genome sequence reads from 827 Watkins landraces, and 217 modern cultivars were mapped to the linearised graph pangenome, and unmapped reads were subsequently assembled. The final assembly resulted in 15,060,124 contigs not in the main graph, with a total length of 8.74 Gb, of which 82.43% was estimated to consist of repetitive sequences, with long terminal repeats (LTRs) being the most prevalent class (61.68%) (Supplementary Table 3). A total of 163,778 gene models were predicted in these contigs, substantially expanding the gene repertoire. Strict functional annotation of gene models identified 34,295 high-confidence annotated genes. Among the landrace AG groups, AG2 had the fewest unique gene models (109), whereas AG6 had the most (518), while 1,114 gene models were shared across all seven AGs (Figure 1; Supplementary Table 4). These results indicate that Watkins landraces harbour substantial additional sequence and gene-content diversity that is absent from modern breeding lines.

**Figure 1.**
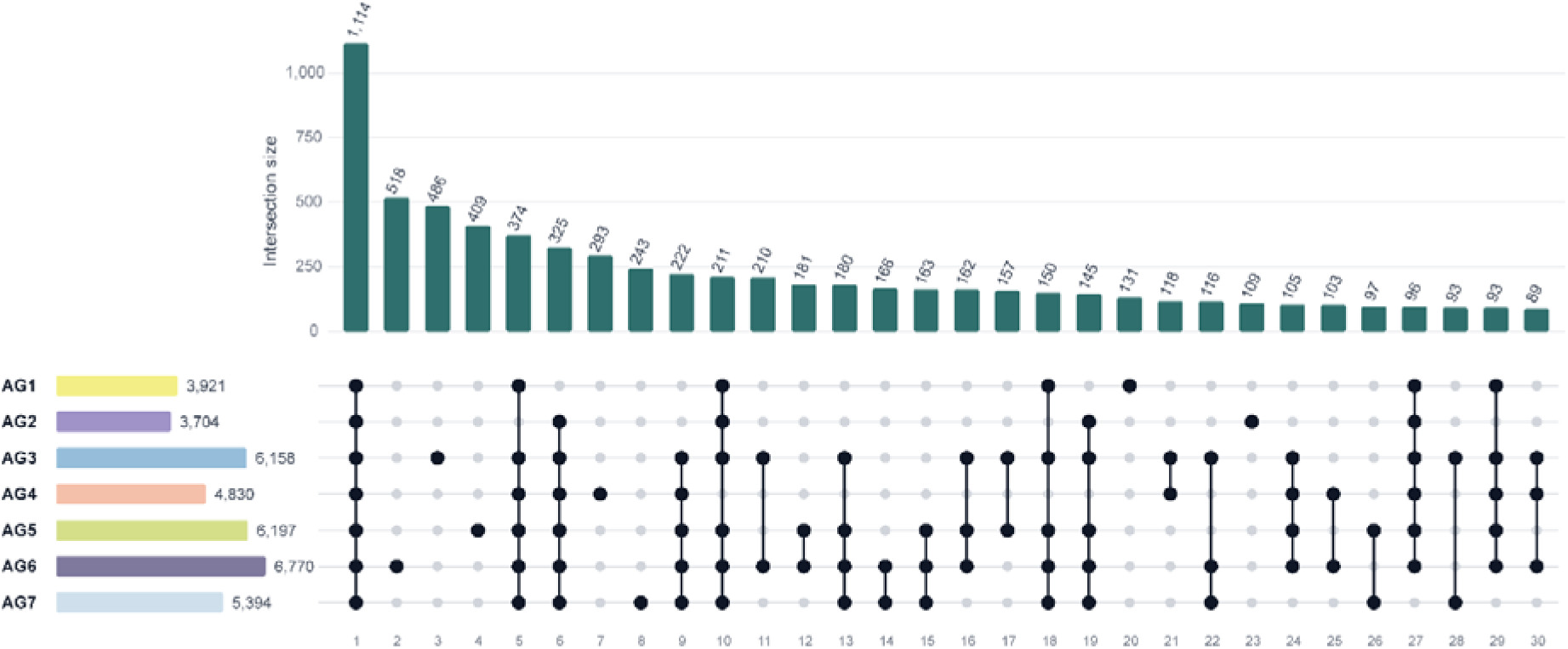
UpSet plot showing the top 30 intersections of Watkins landrace-only gene models across ancestral groups. Horizontal bars (left) show total number of landrace-only gene models present in each AG, summed across all intersections. Vertical bars (top) show the number of gene models exclusive to each intersection. Gene models are considered exclusive to an intersection when PAV data flagged the model as “present” in at least one accession from each AG included within the intersection and “absent” across all accessions of the remaining AGs.

### Comparison with recent wheat pangenomic resources

Wheat pangenomic resources have expanded the representation of sequence diversity beyond a single reference genome. Montenegro et al. (2017) constructed a hexaploid bread wheat pangenome using short-read data from 18 wheat cultivars, recovering approximately 350 Mb of additional sequence and demonstrating extensive gene presence–absence variation (Montenegro *et al*., 2017). Bayer et al. (2022) subsequently developed Wheat Panache, integrating 16 genome assemblies into a 15.8 Gb graph pangenome and providing an interactive resource for exploring genomic regions and gene PAV (Bayer *et al*., 2022). Cheng et al. (2025) constructed an approximately 16.15 Gb graph pangenome from 22 wheat assemblies and identified 140,261 distinct gene families across 19 annotated genomes (Cheng et al., 2025) using OrthoFinder (Emms and Kelly, 2019). In contrast, our graph pangenome comprises 16.44 Gb of sequence derived from 21 genome assemblies, including 1.89 Gb (11.51%) of non-reference sequence, and contains 118,038 gene models. However, these gene numbers are not directly comparable because Cheng et al. (2025) use OrthoFinder to identify gene families rather than characterise the presence or absence of individual genes, as performed in our study.

Besides, White et al. (2025) generated de novo annotations for nine wheat cultivars using genomic assemblies, RNA-seq, Iso-Seq and protein evidence, and investigated variation in gene content and expression across tissues (White et al., 2025). We assessed transcriptional support using 28 publicly available Iso-Seq libraries from 14 wheat cultivars, as reported in White et al. (2025), and aligned the full-length transcripts against our graph pangenome and additional contigs. Only the top hit alignments with ≥95% sequence identity and ≥90% query coverage were retained, 69,060 (58.51%) graph gene models and 3,862 (2.36%) additional gene models were supported by at least one Iso-Seq transcript. Of the 34,295 high-confidence annotated genes, 2,315 (6.75%) showed independent Iso-Seq support. Notably, when the same Iso-Seq transcripts were aligned to their corresponding source genomes using the same alignment criteria, an average of only 32.43% of their annotated genes per cultivar were supported by at least one Iso-Seq transcript (median, 32.73%; range, 29.98–36.33%; Supplementary Table 5), indicating that the relatively low proportion of genes with Iso-Seq support is largely constrained by the available transcriptomic sampling (e.g., limited representation of cultivars and tissues). We further assessed the consistency between Iso-Seq evidence and our gene PAV calls. Across the 14 cultivars, 523,175 graph gene models were supported by Iso-Seq data in their respective source cultivars, of which 504,327 were also called as present in the same cultivar by our PAV analysis, representing an overall concordance of 96.40% (Supplementary Table 5). These results provide independent transcriptional evidence for our gene predictions and support the reliability of our gene PAV analysis.

### Presence/absence variation (PAV) analysis

To assess the extent of novel genomic content present in landraces, we constructed an accumulation curve in two stages, iteratively adding modern cultivars first, followed by landraces. The addition of landraces showed an increase in variable gene model content above the background of genes present in modern lines (Figure 2A and 2B). From the 34,295 new functionally annotated gene models, 668 were exclusive to modern cultivars, and 10,426 were exclusive to landraces, which represent an important reservoir of previously untapped diversity for wheat breeding (Figure 2A). In addition, for all predicted gene models, Watkins landraces contributed a total of 52,236 additional gene models (Figure 2B). Across the 1,044 accessions, the presence frequency distribution of the 34,295 new functionally annotated genes was skewed towards low-frequency variation: 24,205 gene models (70.6%) were present in <15% of accessions and only 74 (0.2%) in ≥99% of accessions (Figure 2C).

**Figure 2.**
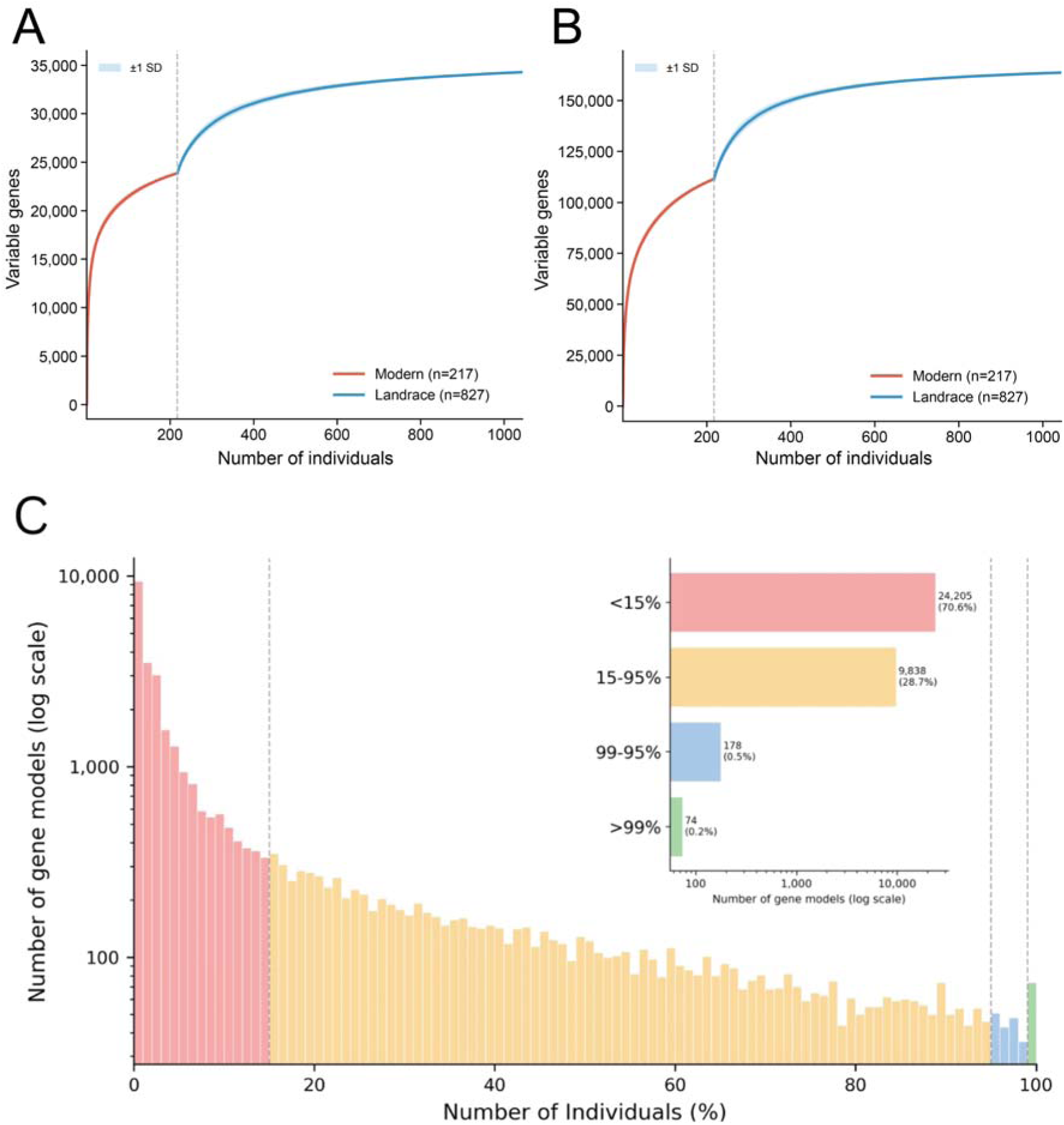
Accumulation curve of variable gene models. (**A**) Accumulation of functionally annotated gene models and (**B**) accumulation of all predicted gene models, with all modern lines added first, followed by Watkins landraces across 100 random permutations. The grey line indicates the addition of the first landrace. (**C**) The presence frequency distribution of 34,295 functionally annotated gene models in the PAV matrix based on the following presence thresholds: >99%, 95–99%, 15-95%, and <15% of individuals. The bar chart indicates the total number of gene models within each category.

Uniform manifold approximation projection (UMAP) (McInnes et al., 2018) visualisation of the 34,295 high-confidence gene models revealed 30 outliers, comprising 29 modern cultivars and one landrace (WATDE0786) (Figure 3A; Figure S2). To determine whether this separation reflected a technical artefact or a biological introgression event, we examined the 581 gene models that were unique to the outlier cluster (Figure S2). We compared their sequences with the genomes of rye (*Secale cereale* Lo7) (Rabanus-Wallace et al., 2021) and bread wheat (*Triticum aestivum* IWGSC) (International Wheat Genome Sequencing Consortium (IWGSC), 2018). The results showed that 393 of 581 gene models matched rye chromosome 1R (Supplementary Table 6). We further aligned the contigs hosting these 581 gene models with the rye genome, which showed that the majority had their best match towards the end of rye chromosome 1R, supporting a 1RS-like origin (Figure S3; Supplementary Table 7). Together, these results indicate that the 30 outliers are likely due to a rye–wheat introgression, consistent with a previously described rye 1RS/1RS.1BL translocation (Rabinovich, 1998). The grouping of WATDE0786 with the 29 modern cultivar outliers is likely due to contamination, mislabelling of samples, or unintended cross-pollination during seed propagation (Quiroz Chavez, 2023).

**Figure 3.**
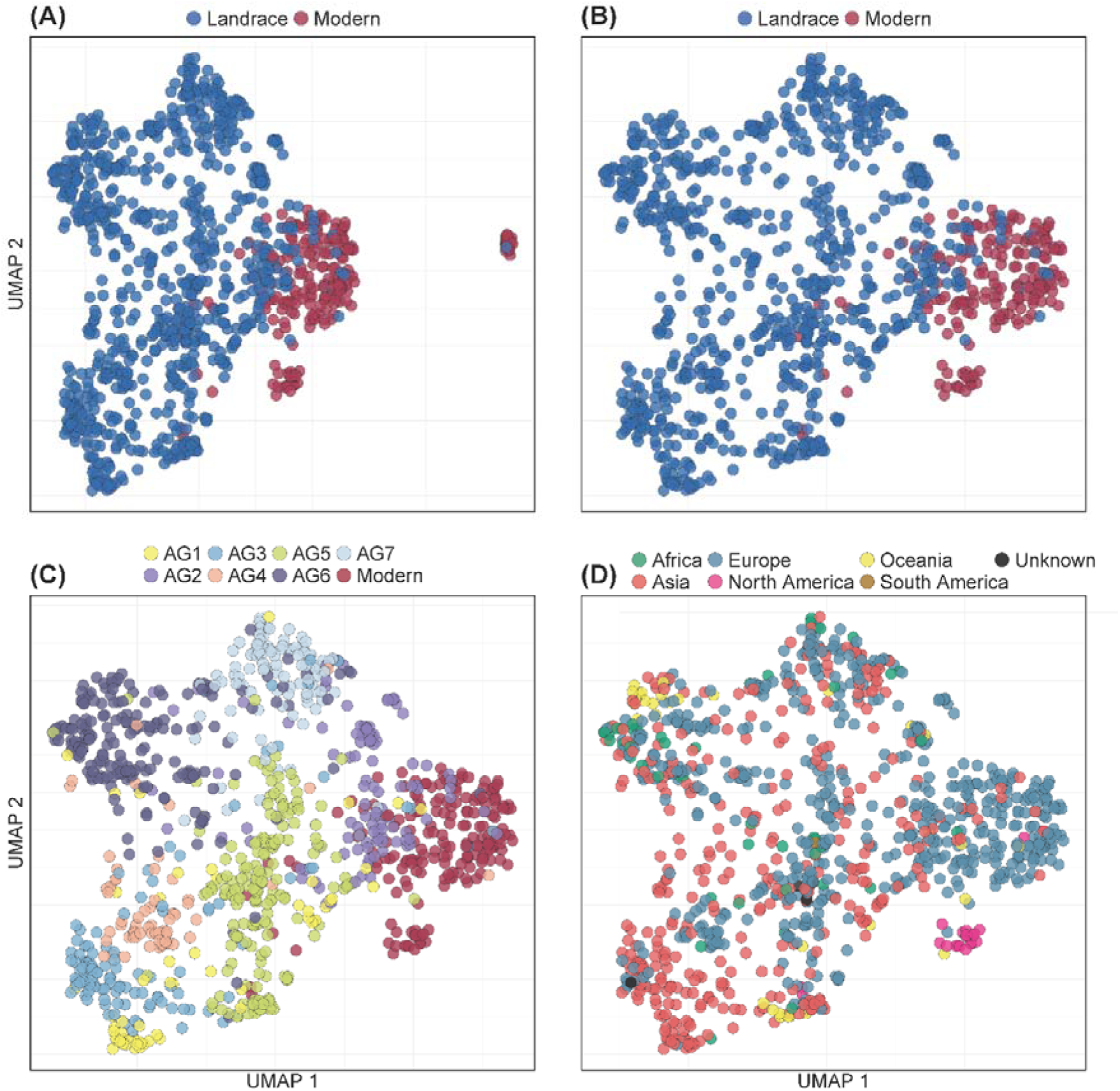
UMAP of the PAV matrix indicates clustering among groups. Uniform manifold approximation projection (UMAP) of the 34,295 functionally annotated gene models from the PAV matrix using the Jaccard distance metric. **(A)** Modern vs landraces with all 1,044 individuals. **(B)** Modern vs landraces, excluding 30 outliers. **(C)** Ancestral groups. **(D)** Continent of origin.

After removing the 30 outliers from the UMAP, the remaining 1,014 individuals clustered between modern cultivars and landrace varieties, with partial overlap between the two groups (Figure 3B). Landraces showed greater dispersion than modern cultivars, which formed a denser and more compact cluster. These results indicate substantially higher genetic diversity among landraces, consistent with their breeding history compared to the intensive breeding of modern cultivars.

Gene model compositional sub-clustering was observed amongst landraces for all AGs, with modern cultivars clustering most closely with AG2 and, to a lesser extent, AG5, while AG1, AG3, AG4, and AG6 showed the greatest separation from modern cultivars (Figure 3C). The pattern is largely consistent with the SNP-based clustering reported by Cheng et al. (2024), indicating that PAV can provide biologically informative signals for population-level analyses (Gordon et al., 2017). Our results support the hypothesis that AG2 and AG5 are the likely founders of modern cultivars. Overlaying the plot with geographic data revealed that UMAP embeddings were also weighted, to a lesser extent, by their source locations (Figure 3D). For example, we observed a small modern-only cluster of individuals obtained almost exclusively from North America, many of which are CIMMYT-derived elite wheat lines, reflecting their shared modern breeding history.

### Phylogenetic analysis based on the gene PAV

To evaluate the phylogenetic signal captured by gene PAV, we constructed a maximum-likelihood tree using the gene PAV matrix (Figure S4). We identified a 277-accession clade dominated by modern cultivars and AG2 accessions, comprising 204 modern cultivars, 70 AG2 individuals, and three additional landraces from AG5 and AG7 (Figure S4). This clade grouped with a 200-accession clade mainly of AG5 individuals (141/200), supporting the close relationship between modern cultivars and AG2/AG5 proposed by Cheng et al. (2024) with SNP-based data (Figure S4). Together, these groups formed a sister lineage to an AG7-majority clade (96/162 individuals), consistent with the reticulate relationships among AG2, AG5, and AG7 inferred by Cheng et al. (2024) (Figure 4; Figure S4). An AG6-majority clade (140/155 individuals) formed a sister group to the AG1+AG3+AG4 clade and was mainly associated with the Iberian Peninsula and Mediterranean origins, whereas the AG1/AG3/AG4 clade was mainly from East, South, and Central/West Asia (Figure 4). This pattern is broadly congruent with the trans-Eurasian dispersal of bread wheat described by Zhao et al. (2023) based on whole-genome sequences of 795 wheat accessions, in which it spread from Southwest Asia towards Europe, South Asia and East Asia. Several AGs formed largely group-specific clades (Figure S4), such as AG6 (140/155 individuals), AG5 (96/118), AG3 (63/72), and AG4 (44/49), which also contributed the largest numbers of AG-specific gene models and UniProt-predicted proteins (Supplementary Table 4). These results indicate that the gene PAV-based phylogeny recapitulates the major ancestry-associated structure inferred from previous SNP-based analyses (Cheng *et al*., 2024; Zhao et al., 2023).

**Figure 4.**
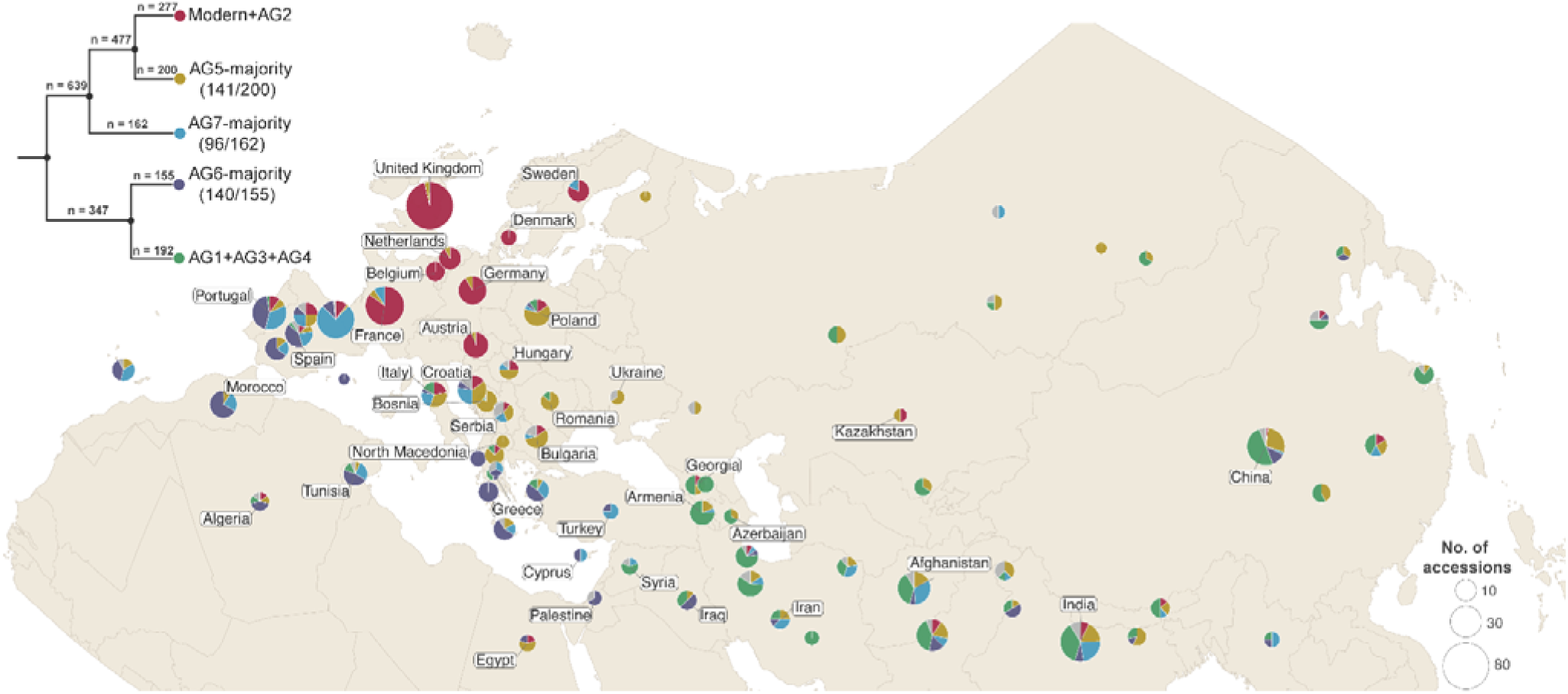
Geographic distribution of major clades inferred from the gene PAV-based phylogeny. Phylogenetic analysis based on gene PAV using Maximum Likelihood inference with 1,000 bootstrap replicates (top left). The accompanying Eurasia-focused map shows the geographical distribution of accessions with geographic origins within the plotted map extent.

### Gene ontology enrichment

The Gene Ontology (GO) enrichment analysis used all functionally annotated hits that were exclusive to landraces as the study set. The background set comprised all functionally annotated genes from the extra contigs, together with all genes from the linearised graph pangenome. GO analysis identified 81 GO terms overrepresented in landraces compared with the background set (enriched, FDR < 0.01) (Figure 5; Supplementary Table 8). Defence response was the most significantly enriched biological process, while response to other organisms and symbiont-induced defence were also enriched. Several terms associated with abiotic stress were also overrepresented, such as response to stress, water deprivation, and oxygen-containing compounds (Figure 5). This suggests that many genes for defence and mycorrhizal interactions may have been lost during the intensive breeding of modern cultivars. In total, 5,772 of the 34,295 high-confidence annotated gene models were predicted to be associated with disease resistance and 4,535 with stress responses. Among the 10,426 landrace-only gene models, 1,423 were associated with disease resistance and 1,322 with stress responses (Supplementary Table 9).

**Figure 5.**
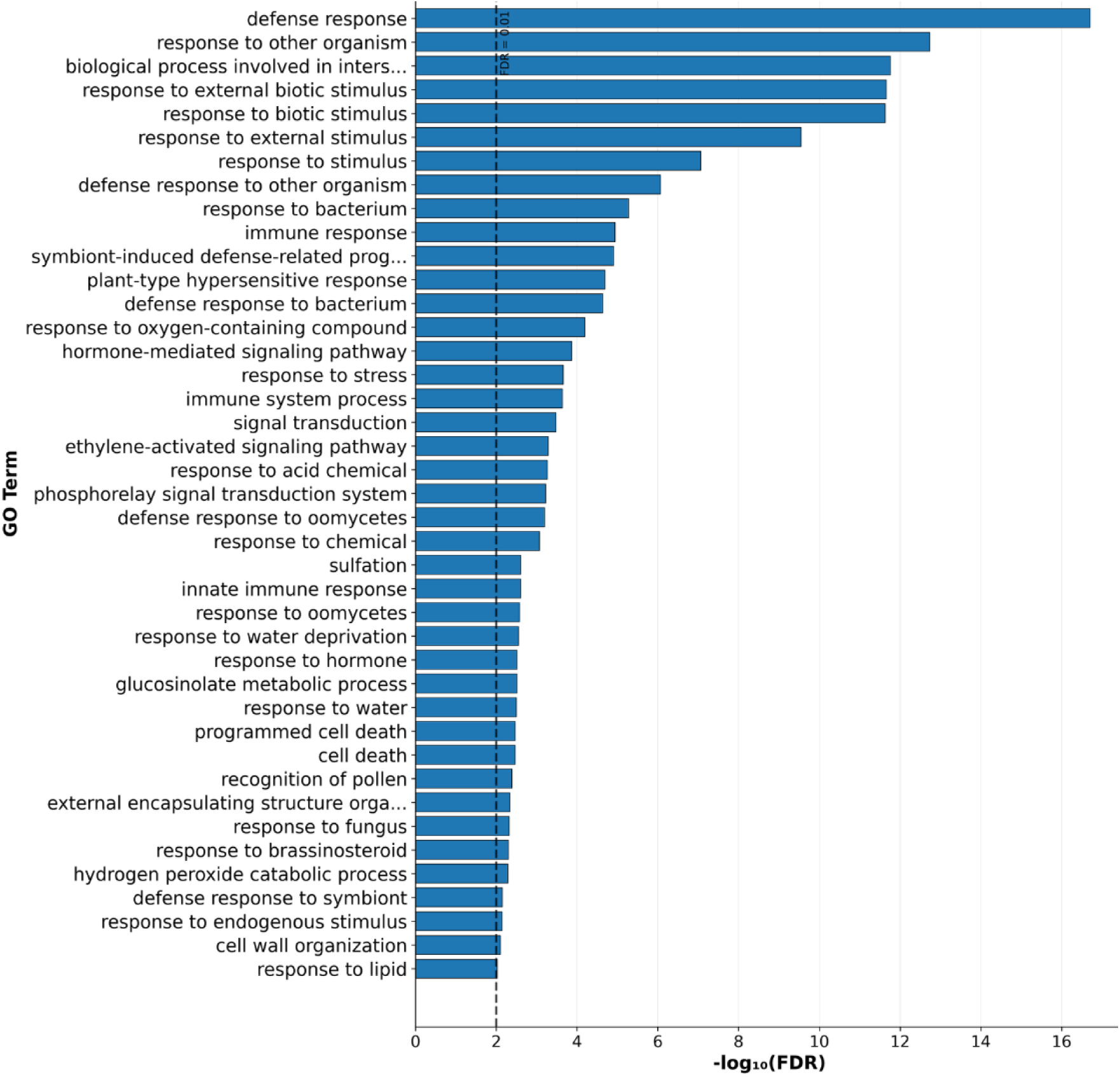
Gene Ontology (GO) term enrichment analysis of variable gene models exclusive to landrace wheat varieties. Bar plot of all significantly enriched GO biological processes ranked by Benjamini-Hochberg False discovery rate (FDR) for landrace individuals.

### NLR gene characterisation

We screened the newly assembled extra contigs for candidate resistance genes encoding NLR motifs using NLR-Annotator (Steuernagel et al., 2020) and identified 1,085 NLR loci, of which 689 were present in all seven landrace ancestry groups and modern cultivars (core) and 176 were specific to the landraces (Figure S5A; Supplementary Table 10). This pattern indicates that wheat landraces retain substantial numbers of resistance-related genes that are absent in modern germplasm. Landrace-specific NLR genes were dominated by the NBARC (Nucleotide-Binding Adaptor shared by APAF-1, R proteins, and CED-4), CC-NBARC (Coiled-Coil-NBARC), and NBARC-LRR (NBARC-Leucine-Rich Repeat) classes, which together accounted for most of the landrace-only NLR repertoire (159 genes; 90.3%) (Figure 6). These genes were most frequently detected in AG4, AG6, and AG3 (Figure S5B), suggesting that multiple AGs contribute to the diversity of NLR-mediated resistance absent from modern wheat lines. The abundance of landrace-specific NLR loci is consistent with a recent genome-wide association study of 297 Watkins accessions, which identified extensive variation in stripe rust resistance across both known resistance regions and potentially novel loci (Singh *et al*., 2026).

**Figure 6.**
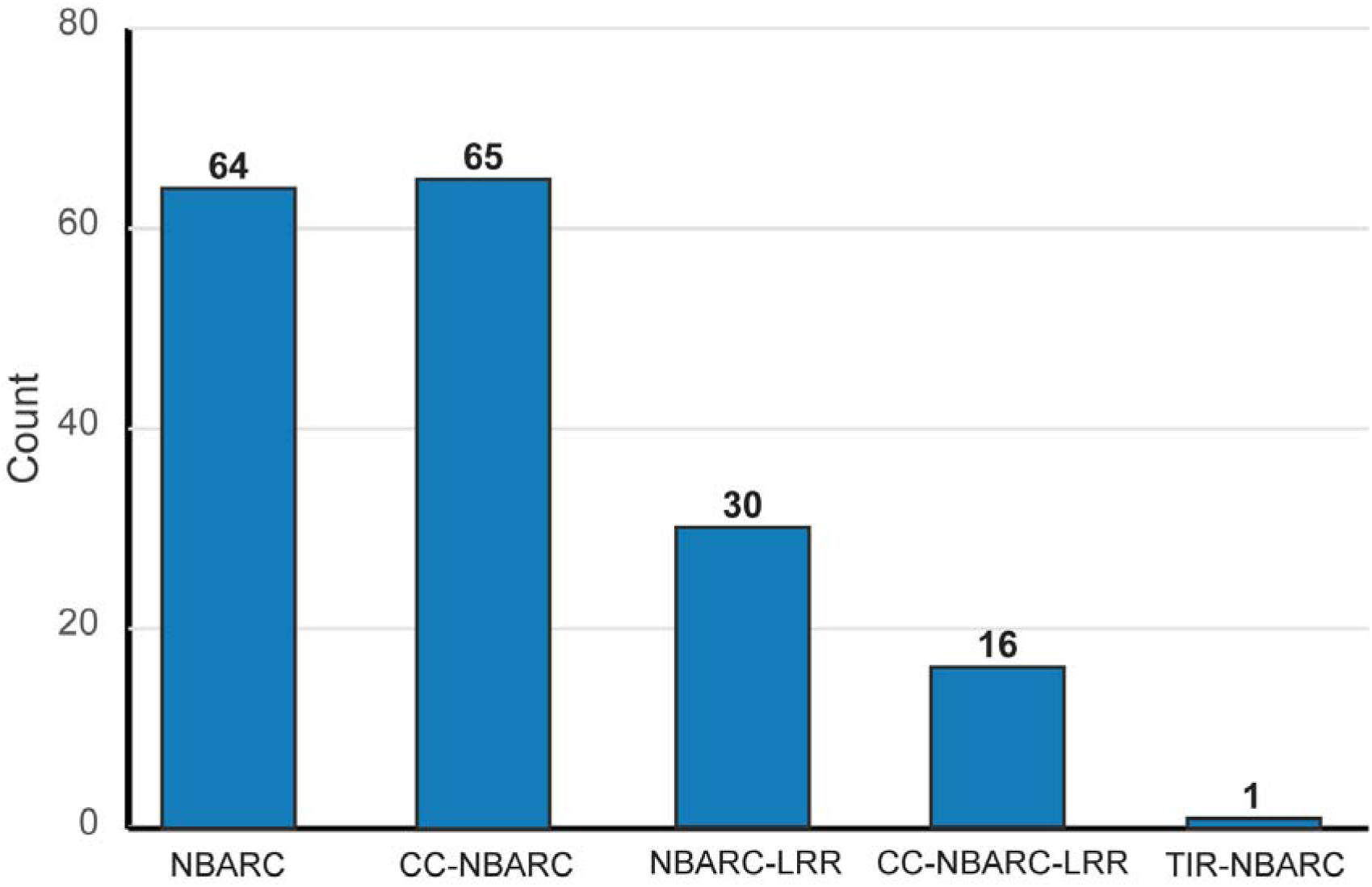
Distribution of NLR structural classes among wheat landraces. Bar plot showing frequency of NLR motifs exclusive to landrace accessions. NBARC (Nucleotide-Binding Adaptor shared by APAF-1, R proteins), CC-NBARC (Coiled-Coil-NBARC), NBARC-LRR (NBARC-Leucine-Rich Repeat), TIR-NBARC (Toll/Interleukin-1 receptor domain NBARC).

To determine whether these NLRs were already represented in the recently published wheat pan-NLRome (Ning et al., 2025), we compared our results with the 2,352 NLR sequences publicly available from the 8,056 NLRs reported by Ning et al. (2025), including 468 classified as landrace-specific. Applying the thresholds that Ning et al. (2025) used to define non-redundant NLRs (≥95% nucleotide identity over ≥90% of the sequence length), we identified 638 of the 2,352 available NLRs (27.1%) in the graph pangenome and a further 107 (4.5%) in the additional contigs, giving a total recovery of 745 sequences (31.7%). Additionally, 78 (16.7%) of the 468 publicly available landrace-specific NLRs from Ning et al. (2025) were recovered in our whole pangenome. In the reciprocal comparison, 171 of the 176 landrace-specific NLR loci identified in our study (97.2%) showed sequence differences to the Ning et al. available NLR sequences. Because only a subset of the 8,056 NLR sequences reported by Ning et al. was publicly accessible at the time of our analysis, we cannot determine whether these loci are also absent from their complete pan-NLRome. Nevertheless, the limited overlap between the currently available datasets suggests that our pangenome-based approach captures additional NLR diversity and reflects differences in NLR identification strategies, as Ning et al. used an approach based on baiting NLR-derived reads followed by accession-specific assembly. Therefore, their targeted approach is inherently enriched for NLR sequences sharing similarity with the known NLR sequences used as bait, whereas our approach identifies novel NLR loci that are not in the genomes used to construct the graph. It should also be noted that the 1,179 NLRs described by Ning et al. (2025) as lost during breeding were defined by significantly reduced frequency in modern cultivars rather than complete absence, and only 641 were reported as landrace-specific. Conversely, the modest recovery of the publicly available Ning et al. sequences in our pangenome suggests that some NLRs, particularly those in highly repetitive regions or present as closely related paralogues, may have been collapsed or removed by our redundancy-filtering thresholds. The two approaches are therefore complementary rather than interchangeable.

To illustrate how these landrace-specific NLRs can be prioritised, we examined a cluster of nine NLR loci that co-occurred in 58 Watkins landraces, mainly from AG4 (18 of 64 accessions), AG3 (15/111) and AG6 (15/193) and originating from Afghanistan, the former USSR, Greece, Iran and Spain, but was absent from all 217 modern cultivars (Supplementary Table 11). The cluster included a complete CC–NBARC–LRR gene (maker-asm3_jcf7180084687145-snap-gene-0.8) encoding a 1,050 amino acid protein, with its best UniProt match corresponding to the rice blast resistance protein Pikm2. Four loci in this cluster were identical or ≥99.7% identical to NLR sequences independently assembled from Watkins landraces and classified as landrace-specific by Ning et al. (2025), providing independent support for their presence in the Watkins germplasm. Read-pair links between the additional contigs and the graph pangenome supported localisation of the cluster to an NLR-rich region at the distal end of chromosome arm 2AL (Chinese Spring RefSeq v1.0, approximately 778.4–778.6 Mb), adjacent to TraesCS2A02G590100, a candidate gene for stripe rust resistance identified by SNP-GWAS of Ning et al. (2025). This highlights a geographically and ancestry-structured landrace-specific NLR cluster that represents a candidate region for functional validation and potential use in wheat pre-breeding.

The breeding value of these landrace-specific genes needs to be considered in the context of how such variation can be deployed. Many successful uses of landrace and wild relative diversity in wheat, such as Sr2 and the 1RS.1BL translocation, involved the transfer of large chromosomal segments rather than individual genes (McFadden, 1930; Rabinovich, 1998), and examples from diploid species such as rice may not translate directly to hexaploid wheat, where homoeologous gene copies can buffer the effect of a single gene. Furthermore, analyses of the Watkins collection have shown that the phenotypic effects of individual loci depend strongly on the genomic background in which they occur (Cheng *et al*., 2024). The presence of a landrace-specific gene therefore does not by itself establish its breeding value. A key feature of the resource presented here is that each gene model is linked to the specific accessions and ancestral groups that carry it, together with the complete gene content of those accessions. This allows candidate genes to be prioritised in donor lines with well-characterised backgrounds, supports the selection of donors for the development of introgression or near-isogenic lines, and enables gene content to be incorporated alongside SNP haplotypes in association and genomic prediction analyses. Functional validation of candidate genes in defined genetic backgrounds, for example using near-isogenic lines, mutant populations or gene editing, will be required to confirm their contribution to stress resilience before deployment in breeding.

To maximise the value of the Watkins wheat pangenome resource and facilitate its use by the wider research and breeding community, we have developed the Watkins Wheat Pangenome Data Explorer (https://pangenome.wheatgenome.info/), an interactive platform that provides public access to the pangenome sequences, gene annotations, functional annotations, and gene PAV. The platform enables users to explore gene presence/absence patterns across ancestral groups, perform BLAST sequence similarity searches against all gene and contig sequences, and interactively visualise the UMAP plot based on gene PAV with accessions highlighted according to selected metadata (e.g., geographic origin).

Together, this study established the largest wheat pangenome assembled to date, capturing gene content variation across more than 1,000 accessions and providing an unprecedented view of the diversity preserved within the Watkins landrace collection. Our analyses reveal that landraces harbour a substantial reservoir of genes absent from modern cultivars, many of which are associated with biotic and abiotic stress responses. These findings indicate that the gene content represented in contemporary wheat breeding captures only a fraction of the diversity present within the species and highlight the extent of functional variation that remains outside elite germplasm. By connecting novel genes to individual accessions and population groups, this resource provides a foundation for dissecting the genetic basis of adaptation and for reintroducing valuable diversity into modern breeding programs. More broadly, the scale of this pangenome demonstrated the power of population-scale analyses to uncover biologically important variation that is inaccessible through single-reference approaches, creating a framework for the systematic exploration and utilisation of wheat genetic diversity.

## Methods

### Graph-based wheat pangenome construction and annotation using 21 varieties

The graph pangenome was constructed using minigraph (Li et al., 2020) v0.21 (options:-f0.1), with the IWGSC Chinese Spring RefSeq v1.0 (International Wheat Genome Sequencing Consortium (IWGSC), 2018) reference genome (GenBank accession number: GCA_900519105.1). The additional 20 genomes included in the graph pangenome were selected based on assembly quality and representation of diverse bread wheat germplasm (Supplementary Table 1). Whole-genome comparison of Chinese Spring RefSeq v1.0 (International Wheat Genome Sequencing Consortium (IWGSC), 2018) with the PacBio HiFi assembly of Fielder (Sato et al., 2021) using minimap2 (Li, 2018) showed overall sequence correspondence between the two assemblies (Figure S1F), supporting the assembly quality of RefSeq v1.0 and its suitability as the backbone for graph construction. The 21 genomes were realigned to the resulting graph using minigraph (Li *et al*., 2020) to infer PAV, and the alignments were converted to BED format using BEDTools v2.30.0 (Quinlan and Hall, 2010). Segment PAVs across the 21 genomes were generated using bedtools multiinter. The segment PAVs were then intersected with publicly available gene annotations for 20 accessions (excluding Attraktion, which lacks genome annotation) (Supplementary Table 1). For each accession, a gene was called present when graph segments classified as present collectively cover at least 75% of the gene’s full annotated length.

### Watkins pangenome assembly and annotation

Sequence data for 827 Watkins landrace cultivars and 217 modern cultivars was downloaded from the National Sequence Archive, China National Centre for Bioinformation (BioProject: PRJCA019636). Four of the modern cultivars in the resequencing dataset were also represented among the 21 genome assemblies used to construct the wheat graph pangenome. Quality control and adapter trimming were performed using Fastp v0.24.0 (Chen, 2023) with default parameters. The final pangenome was assembled using an iterative mapping and assembly approach (Hu et al., 2024; Hu et al., 2020) with the linearised wheat graph as the reference. Batches of clean reads were mapped to the linearised graph pangenome using Bowtie2 (--end-to-end --sensitive) (Langmead and Salzberg, 2012) (Supplementary Table 12). Unmapped reads and singleton reads, together with their read pair, were extracted with Samtools v1.15 (Li et al., 2009), followed by retrieving untrimmed reads using Seqkit v2.10.0 (Shen et al., 2024), as required by MaSuRCA v4.1.2 (Zimin et al., 2013) (cgwErrorRate = 0.15, JF_SIZE = 20,000,000,000) for subsequent *de novo* assembly. Contigs were added to the growing pangenome, and the process was repeated until all read pair batches were processed. Because unmapped reads were pooled across all accessions within each batch prior to assembly, sequences shared by multiple accessions were assembled from their combined coverage rather than from the ∼12.7× coverage of an individual accession. This iterative mapping and assembly approach has previously been applied to hexaploid wheat (Montenegro *et al*., 2017) and other crops (Hu *et al*., 2024; Hu *et al*., 2020), and resequencing depths of ∼10× have been shown to be sufficient for reliable gene presence/absence calling (Hu *et al*., 2025). Gene models present in only one or a few accessions are assembled from correspondingly lower coverage and may therefore be under-represented. Contigs shorter than 200 bp were discarded. The remaining contigs were compared with the NCBI core-nt database (29 May 2025) using BLAST v2.16.0 (Camacho et al., 2009) (blastn-task dc-megablast-evalue 1e-5). Contigs with the best hit to non-green plant, chloroplast, or mitochondrial sequences were excluded from further analysis. Redundant contigs were removed by (i) aligning the newly assembled contigs with the linearised graph pangenome using NCBI-BLAST+ (Camacho *et al*., 2009) and removing contigs with ≥95% sequence similarity and ≥90% coverage; the BLASTN alignments were further evaluated at the query-subject pair level by merging non-overlapping query intervals from HSPs with ≥99% identity, and contigs with ≥99% combined query coverage were removed (ii) aligning the contigs against the 21 genomes used to construct the graph pangenome using minimap2 (Li, 2018) and applying the same thresholds (≥95% identity and ≥90% coverage) to remove redundant contigs, (iii) aligning the remaining contigs with themselves with minimap2 (Li, 2018) to eliminate redundancy using the threshold of ≥99% similarity and ≥95% sequence coverage, and (iv) aligning the retained contigs back to the graph pangenome using minigraph (Li *et al*., 2020). Contigs with ≥95% sequence identity and ≥90% coverage were removed.

To identify transposable elements (TEs), two curated TE libraries of TREP (Wicker et al., 2002) and ClariTeRep (https://github.com/jdaron/CLARI-TE) were combined and used as the input library for the Extensive de novo TE Annotator (EDTA) v.2.1.0 (Ou et al., 2019) with default parameters. Newly identified elements were merged with the curated repeat sequences and subsequently used to mask repetitive elements for all contigs using RepeatMasker v4.1.0 (Tarailo-Graovac and Chen, 2009). *De novo* annotation was performed with three rounds of MAKER2 v2.31.9 (Holt and Yandell, 2011), the first round incorporated protein and EST evidence from the Chinese Spring IWGSC RefSeq v2.1 assembly (Zhu et al., 2021), while the second and third rounds incorporated *ab initio* gene predictions generated by SNAP (ZOE library version 2013-02-16) (Korf, 2004) and AUGUSTUS v3.3.3 (Stanke and Waack, 2003).

### Gene presence/absence variation analysis

For each accession, sequencing reads that did not map to the reference genome were aligned to the contigs generated via *de novo* assembly. The GFF file for these contigs was converted to BED format for compatibility with downstream tools. Mosdepth was used to quantify per-gene read depth across 163,778 predicted gene models with the --fast_mode flag (v0.3.13) (Pedersen and Quinlan, 2018). A gene model was classified as “present” (1) in an accession when at least 95% of its length was covered by one or more sequencing reads; otherwise, it was classified as ‘absent’ (0). Gene models failing to meet this threshold were flagged as “absent” (0), following the approach used by the SGSgeneloss (Golicz et al., 2015). Gene models flagged as absent across all genomes were removed from further analysis.

Protein sequences for the predicted gene models were extracted with gffread (v0.12.7) (Pertea and Pertea, 2020). These were compared with the UniProt/SwissProt database (v2026_01, The UniProt Consortium) (UniProt Consortium, 2025) using MMseqs2 (v18.8cc5c) (Steinegger and Söding, 2017) with parameterse 1e-3,-s 7.5,-max-seqs 1. Hits were classified as “high confidence” if eval < 0.001. For gene models with multiple high-confidence hits, the hit with the lowest e-value was retained. The gene-model PAV matrix was filtered to retain only rows that returned high-confidence hits and was used for all downstream analyses.

Variable gene model accumulation curves were constructed using 100 random permutations. The mean and standard deviation of gene model count for each variable were calculated as new genomes were added.

Uniform manifold approximation and projection (UMAP) was applied to the subset PAV matrix using the umap-learn Python package (v0.5) (McInnes *et al*., 2018). UMAP Embeddings were calculated using the Jaccard distance metric. Embeddings were generated and calculated using the following parameters: n_neighbors=50, min_dist=0.1, spread=1.0. Plots were generated with Python v3.11, R v4.5.2 (R Core Team, 2026), and Matplotlib (v3.10.8) (Hunter, 2007).

### Iso-Seq validation of gene predictions and gene PAV

To independently assess transcriptional support for the predicted gene models, we downloaded 28 publicly available PacBio Iso-Seq libraries from the European Nucleotide Archive (PRJEB51827) representing 14 wheat cultivars, as reported by White et al. (2025). Full-length Iso-Seq transcripts were aligned against the linearised graph pangenome and the additional contigs using minimap2 with the parameters-x splice:hq. For each transcript, only the top-scoring alignment with ≥95% sequence identity and ≥ 90% query coverage was retained. For each cultivar, supported graph models were compared with the corresponding cultivar’s calls in the gene PAV matrix, and concordance was calculated as the proportion called present. To evaluate the extent of Iso-Seq support in the source cultivars, the same Iso-Seq reads were also aligned separately to their corresponding source genomes using the same alignment criteria. 14 cultivar annotations were obtained from Ensembl Plants release 52 for the nine cultivars annotated by White et al. (2025) and release 59 for the five additional cultivars.

### Gene ontology enrichment

Gene ontology terms were obtained using the GO term mapping file from Uniprot (v 2025.04). GO: IDs were mapped to human-readable descriptions using go-basic.obo. Gene enrichment was applied using the Goatools Python package (v1.4.12) GoEnrichmentStudy() function, with parameters: alpha=0.01, methods=[fdr_bh], propagate_counts=True. Fisher’s exact test was used to assess significance, with FDR correction. Overrepresented GO terms that passed FDR correction with a p-value < 0.01 were considered enriched. The significant GO terms were then plotted with Python 3.12. Those annotated with “defense response” (GO:0006952) or any of its descendant terms were considered associated with disease resistance; those annotated with “response to stress” (GO:0006950) or its descendants, excluding terms under defense response, response to biotic stimulus (GO:0009607) and immune system process (GO:0002376), were considered associated with stress responses.

### NLR gene identification

NLR immune receptor loci were identified directly from the additional contigs using NLR-Annotator (Steuernagel *et al*., 2020) (v2). NLR-Annotator output files were used for all downstream NLR analyses. NLR predictions were converted back to their genomic coordinates on the original contigs, with overlapping predictions on the same contig and strand considered duplicate representations of the same NLR locus and collapsed by retaining the NLR loci with the longer interval.

To compare our NLR predictions with the recently reported wheat pan-NLRome of Ning et al. (2025), we found that the complete set of 8,056 NLR sequences deposited in the China National Genomics Data Center under BioProject PRJCA047178 was not publicly available at the time of our analysis; however, we were able to retrieve 2,352 NLR sequences from GenBase, including 468 classified as landrace-specific. These sequences were aligned to our graph using minigraph (Li *et al*., 2020) v0.21 (-c-x lr) and to the additional contigs with minimap2 (Li, 2018) (-x asm20-k17). A sequence was considered represented in our pangenome when an alignment had ≥95% sequence identity and ≥90% query coverage.

## Data availability

The previously published whole-genome resequencing data used in this study are publicly available from the National Sequence Archive of the China National Center for Bioinformation under BioProject accession PRJCA019636. The pangenome sequences, gene annotations, functional annotations, and gene presence–absence variation matrix are accessible through the Watkins Wheat Pangenome Data Explorer (https://pangenome.wheatgenome.info/).

## Author contributions

**T. L.:** Writing original draft, Formal analysis, Methodology, Investigation, Visualisation. **J. Br.:** Methodology, Investigation, Writing original draft, Visualisation, Website development. **F. E. A.:** Investigation, Writing original draft, Methodology, Visualisation. **M. B.:** Data curation, Formal analysis, Methodology. **A. D.:** Formal analysis, Methodology, Visualisation. **T. M.:** Investigation. **T. B.:** Investigation. **P. B.:** Data curation, Investigation, Formal analysis, Methodology. **J. Ba.:** Conceptualisation, Funding acquisition. **D. E.:** Conceptualisation, Funding acquisition, Writing – review and editing.

## Acknowledgements

This work was supported by resources provided by the Pawsey Supercomputing Research Centre’s Setonix Supercomputer (https://doi.org/10.48569/18sb-8s43), with funding from the Australian Government and the Government of Western Australia. We acknowledge funding from the Australian Research council projects DP200100762, DP210100296, FL230100030 and LP230100351. We acknowledge the use of New Zealand eScience Infrastructure (NeSI) high-performance computing facilities. The authors declare no conflict of interest.

## References

Amas, J.C., Bayer, P.E., Hong Tan, W., Tirnaz, S., Thomas, W.J.W., Edwards, D., and Batley, J. (2023). Comparative pangenome analyses provide insights into the evolution of *Brassica rapa* resistance gene analogues (RGAs). Plant Biotechnol J 21:2100–2112. 10.1111/pbi.14116.

Anderson, R., Bayer, P.E., and Edwards, D. (2020). Climate change and the need for agricultural adaptation. Curr Opin Plant Biol 56:197–202. 10.1016/j.pbi.2019.12.006.

Barbosa, P.A.M., Fritsche-Neto, R., Andrade, M.C., Petroli, C.D., Burgueno, J., Galli, G., Willcox, M.C., Sonder, K., Vidal-Martinez, V.A., Sifuentes-Ibarra, E., et al. (2021). Introgression of maize diversity for drought tolerance: subtropical maize landraces as source of new positive variants. Front Plant Sci 12:691211. 10.3389/fpls.2021.691211.

Bayer, P.E., Golicz, A.A., Scheben, A., Batley, J., and Edwards, D. (2020). Plant pan-genomes are the new reference. Nature Plants 6:914–920. 10.1038/s41477-020-0733-0.

Bayer, P.E., Petereit, J., Durant, E., Monat, C., Rouard, M., Hu, H., Chapman, B., Li, C., Cheng, S., Batley, J., et al. (2022). Wheat Panache: A pangenome graph database representing presence-absence variation across sixteen bread wheat genomes. Plant Genome 15:e20221. 10.1002/tpg2.20221.

Blower, A., Ray, R.V., Rawsthorne, S., Howell, P.J., Leigh, F.J., and Kanyuka, K. (2025). Harnessing primary, secondary and tertiary genepools for durable wheat disease resistance. Theor Appl Genet 138:270. 10.1007/s00122-025-05053-0.

Camacho, C., Coulouris, G., Avagyan, V., Ma, N., Papadopoulos, J., Bealer, K., and Madden, T.L. (2009). BLAST+: architecture and applications. BMC Bioinformatics 10:421. 10.1186/1471-2105-10-421.

Chen, S. (2023). Ultrafast one-pass FASTQ data preprocessing, quality control, and deduplication using fastp. iMeta 2:e107. 10.1002/imt2.107.

Cheng, H., Kong, L., Zhu, K., Zhao, H., Li, X., Zhang, Y., Ning, W., Jiang, M., Song, B., and Cheng, S. (2025). Structural variation-based and gene-based pangenome construction reveals untapped diversity of hexaploid wheat. J Genet Genomics 52:774–785. 10.1016/j.jgg.2025.03.015.

Cheng, S., Feng, C., Wingen, L.U., Cheng, H., Riche, A.B., Jiang, M., Leverington-Waite, M., Huang, Z., Collier, S., Orford, S., et al. (2024). Harnessing landrace diversity empowers wheat breeding. Nature 632:823–831. 10.1038/s41586-024-07682-9.

Danilevicz, M.F., Fernandez, C.G.T., Marsh, J.I., Bayer, P.E., and Edwards, D. (2020). Plant pangenomics: approaches, applications and advancements. Curr Opin Plant Biol 54:18–25. 10.1016/j.pbi.2019.12.005.

Dwivedi, S.L., Ceccarelli, S., Blair, M.W., Upadhyaya, H.D., Are, A.K., and Ortiz, R. (2016). Landrace Germplasm for Improving Yield and Abiotic Stress Adaptation. Trends Plant Sci 21:31–42. 10.1016/j.tplants.2015.10.012.

Emms, D.M., and Kelly, S. (2019). OrthoFinder: phylogenetic orthology inference for comparative genomics. Genome Biol 20:238. 10.1186/s13059-019-1832-y.

Golicz, A.A., Batley, J., and Edwards, D. (2016). Towards plant pangenomics. Plant Biotechnol J 14:1099–1105. 10.1111/pbi.12499.

Golicz, A.A., Martinez, P.A., Zander, M., Patel, D.A., Van De Wouw, A.P., Visendi, P., Fitzgerald, T.L., Edwards, D., and Batley, J. (2015). Gene loss in the fungal canola pathogen *Leptosphaeria maculans*. Funct Integr Genomics 15:189–196. 10.1007/s10142-014-0412-1.

Gordon, S.P., Contreras-Moreira, B., Woods, D.P., Des Marais, D.L., Burgess, D., Shu, S., Stritt, C., Roulin, A.C., Schackwitz, W., Tyler, L., et al. (2017). Extensive gene content variation in the *Brachypodium distachyon* pan-genome correlates with population structure. Nat Commun 8:2184. 10.1038/s41467-017-02292-8.

Guo, D., Li, Y., Lu, H., Zhao, Y., Kurata, N., Wei, X., Wang, A., Wang, Y., Zhan, Q., Fan, D., et al. (2025). A pangenome reference of wild and cultivated rice. Nature 642:662–671. 10.1038/s41586-025-08883-6.

Guo, N., Wang, S., Wang, T., Duan, M., Zong, M., Miao, L., Han, S., Wang, G., Liu, X., Zhang, D., et al. (2024). A graph-based pan-genome of *Brassica oleracea* provides new insights into its domestication and morphotype diversification. Plant Commun 5:100791. 10.1016/j.xplc.2023.100791.

Hao, M., Zhang, L., Ning, S., Huang, L., Yuan, Z., Wu, B., Yan, Z., Dai, S., Jiang, B., Zheng, Y., et al. (2020). The resurgence of introgression breeding, as exemplified in wheat improvement. Front Plant Sci 11:252. 10.3389/fpls.2020.00252.

Haudry, A., Cenci, A., Ravel, C., Bataillon, T., Brunel, D., Poncet, C., Hochu, I., Poirier, S., Santoni, S., Glemin, S., et al. (2007). Grinding up wheat: a massive loss of nucleotide diversity since domestication. Mol Biol Evol 24:1506–1517. 10.1093/molbev/msm077.

Holt, C., and Yandell, M. (2011). MAKER2: an annotation pipeline and genome-database management tool for second-generation genome projects. BMC Bioinformatics 12:491. 10.1186/1471-2105-12-491.

Hu, H., Li, R., Zhao, J., Batley, J., and Edwards, D. (2024). Technological development and advances for constructing and analyzing plant pangenomes. Genome Biol Evol 16:evae081. 10.1093/gbe/evae081.

Hu, H., Zhao, J., Thomas, W.J.W., Batley, J., and Edwards, D. (2025). The role of pangenomics in orphan crop improvement. Nat Commun 16:118. 10.1038/s41467-024-55260-4.

Hu, H., Yuan, Y., Bayer, P.E., Fernandez, C.T., Scheben, A., Golicz, A.A., and Edwards, D. (2020). Legume pangenome construction using an iterative mapping and assembly approach. In Legume Genomics: Methods and Protocols, M. Jain and R. Garg, eds. (Springer US: New York, NY), pp. 35–47. 10.1007/978-1-0716-0235-5_3.

Hunter, J.D. (2007). Matplotlib: a 2D graphics environment. Comput Sci Eng 9:90–95. 10.1109/MCSE.2007.55.

International Wheat Genome Sequencing Consortium (IWGSC) (2018). Shifting the limits in wheat research and breeding using a fully annotated reference genome. Science 361:eaar7191. 10.1126/science.aar7191.

Jiao, C., Xie, X., Hao, C., Chen, L., Xie, Y., Garg, V., Zhao, L., Wang, Z., Zhang, Y., Li, T., et al. (2025). Pan-genome bridges wheat structural variations with habitat and breeding. Nature 637:384–393. 10.1038/s41586-024-08277-0.

Korf, I. (2004). Gene finding in novel genomes. BMC Bioinformatics 5:59. 10.1186/1471-2105-5-59.

Langmead, B., and Salzberg, S.L. (2012). Fast gapped-read alignment with Bowtie 2. Nat Methods 9:357–359. 10.1038/nmeth.1923.

Li, H. (2018). Minimap2: pairwise alignment for nucleotide sequences. Bioinformatics 34:3094–3100. 10.1093/bioinformatics/bty191.

Li, H., Feng, X., and Chu, C. (2020). The design and construction of reference pangenome graphs with minigraph. Genome Biol 21:265. 10.1186/s13059-020-02168-z.

Li, H., Handsaker, B., Wysoker, A., Fennell, T., Ruan, J., Homer, N., Marth, G., Abecasis, G., Durbin, R., and Genome Project Data Processing, S. (2009). The Sequence Alignment/Map format and SAMtools. Bioinformatics 25:2078–2079. 10.1093/bioinformatics/btp352.

Li, T., Mohamedikbal, S., Bestry, M., Batley, J., and Edwards, D. (2025). Pangenomics combined with artificial intelligence and precision breeding can accelerate crop improvement. Curr Opin Plant Biol 88:102825. 10.1016/j.pbi.2025.102825.

Liu, Q., Zhang, X., Su, Y.H., and Zhang, X.S. (2022). Genetic mechanisms of cold signaling in wheat (*Triticum aestivum* L.). Life (Basel) 12:700. 10.3390/life12050700.

Lobell, D.B., Schlenker, W., and Costa-Roberts, J. (2011). Climate trends and global crop production since 1980. Science 333:616–620. doi:10.1126/science.1204531.

Ma, W., Liu, Y., Wei, X., Zhang, X., Li, X., Liu, Z., Yuan, L., Li, G., Zhang, S., Yang, Q., et al. (2026). Gapless pangenome analyses reveal fast *Brassica rapa* subspeciation. Science 391:eady7590. 10.1126/science.ady7590.

MacNish, T.R., Kopalli, V., Zanini, S.F., Snowdon, R.J., Golicz, A.A., and Edwards, D. (2026). Accessing crop genetic diversity via pangenomics. Theor Appl Genet 139:96. 10.1007/s00122-026-05201-0.

McFadden, E.S. (1930). A successful transfer of emmer characters to vulgare wheat. Agron J 22:1020–1034.

McInnes, L., Healy, J., and Melville, J. (2018). Umap: Uniform manifold approximation and projection for dimension reduction. arXiv preprint arXiv:1802.03426.

Montenegro, J.D., Golicz, A.A., Bayer, P.E., Hurgobin, B., Lee, H., Chan, C.K., Visendi, P., Lai, K., Dolezel, J., Batley, J., et al. (2017). The pangenome of hexaploid bread wheat. Plant J 90:1007–1013. 10.1111/tpj.13515.

Neeraja, C.N., Maghirang-Rodriguez, R., Pamplona, A., Heuer, S., Collard, B.C., Septiningsih, E.M., Vergara, G., Sanchez, D., Xu, K., Ismail, A.M., et al. (2007). A marker-assisted backcross approach for developing submergence-tolerant rice cultivars. Theor Appl Genet 115:767–776. 10.1007/s00122-007-0607-0.

Ning, W., Ye, Z., Ye, S., Xian, W., Zou, T., Liu, Z., Wang, W., Chen, B., Xie, W., Cheng, S., et al. (2025). The pan-NLRome and diversity of Watkins wheat provide genetic resources for improving disease resistance and adaptation. Plant Commun 6:101563. 10.1016/j.xplc.2025.101563.

Ou, S., Su, W., Liao, Y., Chougule, K., Agda, J.R.A., Hellinga, A.J., Lugo, C.S.B., Elliott, T.A., Ware, D., Peterson, T., et al. (2019). Benchmarking transposable element annotation methods for creation of a streamlined, comprehensive pipeline. Genome Biol 20:275. 10.1186/s13059-019-1905-y.

Pedersen, B.S., and Quinlan, A.R. (2018). Mosdepth: quick coverage calculation for genomes and exomes. Bioinformatics 34:867–868. 10.1093/bioinformatics/btx699.

Pertea, G., and Pertea, M. (2020). GFF Utilities: GffRead and GffCompare. F1000Res 9:304. 10.12688/f1000research.23297.2.

Placido, D.F., Campbell, M.T., Folsom, J.J., Cui, X., Kruger, G.R., Baenziger, P.S., and Walia, H. (2013). Introgression of novel traits from a wild wheat relative improves drought adaptation in wheat. Plant Physiol 161:1806–1819. 10.1104/pp.113.214262.

Qiao, W., Qi, L., Cheng, Z., Su, L., Li, J., Sun, Y., Ren, J., Zheng, X., and Yang, Q. (2016). Development and characterization of chromosome segment substitution lines derived from *Oryza rufipogon* in the genetic background of *O. sativa spp. indica* cultivar 9311. BMC Genomics 17:580. 10.1186/s12864-016-2987-5.

Quinlan, A.R., and Hall, I.M. (2010). BEDTools: a flexible suite of utilities for comparing genomic features. Bioinformatics 26:841–842. 10.1093/bioinformatics/btq033.

Quiroz Chavez, J. (2023). Wheat haplotype diversity by a k-mer based approach. University of East Anglia.

R Core Team (2026). R: a language and environment for statistical computing. https://www.R-project.org/.

Rabanus-Wallace, M.T., Hackauf, B., Mascher, M., Lux, T., Wicker, T., Gundlach, H., Baez, M., Houben, A., Mayer, K.F.X., Guo, L., et al. (2021). Chromosome-scale genome assembly provides insights into rye biology, evolution and agronomic potential. Nat Genet 53:564–573. 10.1038/s41588-021-00807-0.

Rabinovich, S.V. (1998). Importance of wheat-rye translocations for breeding modern cultivar of Triticum aestivum L. Euphytica 100:323–340.

Reif, J.C., Zhang, P., Dreisigacker, S., Warburton, M.L., van Ginkel, M., Hoisington, D., Bohn, M., and Melchinger, A.E. (2005). Wheat genetic diversity trends during domestication and breeding. Theor Appl Genet 110:859–864. 10.1007/s00122-004-1881-8.

Roy, N., Islam, S., Yu, Z.T., Lu, M.Q., Lafiandra, D., Zhao, Y., Anwar, M., Mayer, J.E., and Ma, W.J. (2020). Introgression of an expressed HMW 1Ay glutenin subunit allele into bread wheat cv. Lincoln increases grain protein content and breadmaking quality without yield penalty. Theor Appl Genet 133:517–528. 10.1007/s00122-019-03483-1.

Sato, K., Abe, F., Mascher, M., Haberer, G., Gundlach, H., Spannagl, M., Shirasawa, K., and Isobe, S. (2021). Chromosome-scale genome assembly of the transformation-amenable common wheat cultivar ‘Fielder’. DNA Res 28:dsab008. 10.1093/dnares/dsab008.

Schreiber, M., Jayakodi, M., Stein, N., and Mascher, M. (2024). Plant pangenomes for crop improvement, biodiversity and evolution. Nat Rev Genet 25:563–577. 10.1038/s41576-024-00691-4.

Shen, W., Sipos, B., and Zhao, L. (2024). SeqKit2: A Swiss army knife for sequence and alignment processing. iMeta 3:e191. 10.1002/imt2.191.

Sibikeev, S.N., Baranova, O.A., and Druzhin, A.E. (2021). A prebreeding study of introgression spring bread wheat lines carrying combinations of stem rust resistance genes, *Sr22*+*Sr25* and *Sr35*+*Sr25*. Vavilovskii Zh Genet 25:713–722. 10.18699/Vj21.081.

Singh, J., Awan, M.J.A., Kumar, N., Holden, S., Khangura, R.S., and Brar, G.S. (2026). Watkins wheat landraces: a treasure of stripe rust resistance alleles identified using multi-model association analyses. Theor Appl Genet 139:190. 10.1007/s00122-026-05291-w.

Singh, V.K., Singh, B.D., Kumar, A., Maurya, S., Krishnan, S.G., Vinod, K.K., Singh, M.P., Ellur, R.K., Bhowmick, P.K., and Singh, A.K. (2018). Marker-assisted introgression of *Saltol* QTL enhances seedling stage salt tolerance in the rice variety “Pusa Basmati 1”. Int J Genomics 2018:8319879. 10.1155/2018/8319879.

Smale, M., Reynolds, M.P., Warburton, M., Skovmand, B., Trethowan, R., Singh, R.P., Ortiz-Monasterio, I., and Crossa, J. (2002). Dimensions of diversity in modern spring bread wheat in developing countries from 1965. Crop Sci 42:1766–1779. 10.2135/cropsci2002.1766.

Stanke, M., and Waack, S. (2003). Gene prediction with a hidden Markov model and a new intron submodel. Bioinformatics 19 Suppl 2:ii215–225. 10.1093/bioinformatics/btg1080.

Steinegger, M., and Söding, J. (2017). MMseqs2 enables sensitive protein sequence searching for the analysis of massive data sets. Nat Biotechnol 35:1026– 1028. 10.1038/nbt.3988.

Steuernagel, B., Witek, K., Krattinger, S.G., Ramirez-Gonzalez, R.H., Schoonbeek, H.J., Yu, G., Baggs, E., Witek, A.I., Yadav, I., Krasileva, K.V., et al. (2020). The NLR-Annotator tool enables annotation of the intracellular immune receptor repertoire. Plant Physiol 183:468–482. 10.1104/pp.19.01273.

Tarailo-Graovac, M., and Chen, N. (2009). Using RepeatMasker to identify repetitive elements in genomic sequences. Current protocols in bioinformatics / editoral board, Andreas D. Baxevanis…[et al.]:4.10.11–14.10.14. 10.1002/0471250953.bi0410s25.

Toit, D.F. (1989). Components of resistance in three bread wheat lines to Russian wheat aphid (Homoptera: Aphididae) in South Africa. J Econ Entomol 82:1779–1781. 10.1093/jee/82.6.1779.

UniProt Consortium (2025). UniProt: the Universal Protein Knowledgebase in 2025. Nucleic Acids Res 53:D609–D617. 10.1093/nar/gkae1010.

Valdez, V.A., Byrne, P.F., Lapitan, N.L.V., Peairs, F.B., Bernardo, A., Bai, G.H., and Haley, S.D. (2012). Inheritance and Genetic Mapping of Russian Wheat Aphid Resistance in Iranian Wheat Landrace Accession PI 626580. Crop Sci 52:676–682. 10.2135/cropsci2011.06.0331.

Walkowiak, S., Gao, L., Monat, C., Haberer, G., Kassa, M.T., Brinton, J., Ramirez-Gonzalez, R.H., Kolodziej, M.C., Delorean, E., Thambugala, D., et al. (2020). Multiple wheat genomes reveal global variation in modern breeding. Nature 588:277–283. 10.1038/s41586-020-2961-x.

White, B., Lux, T., Rusholme-Pilcher, R., Juhasz, A., Kaithakottil, G., Duncan, S., Simmonds, J., Rees, H., Wright, J., Colmer, J., et al. (2025). De novo annotation reveals transcriptomic complexity across the hexaploid wheat pan-genome. Nat Commun 16:8538. 10.1038/s41467-025-64046-1.

Wicker, T., Matthews, D.E., and Keller, B. (2002). TREP: a database for Triticeae repetitive elements. Trends Plant Sci 7:561–562. 10.1016/S1360-1385(02)02372-5.

Winfield, M.O., Allen, A.M., Wilkinson, P.A., Burridge, A.J., Barker, G.L.A., Coghill, J., Waterfall, C., Wingen, L.U., Griffiths, S., and Edwards, K.J. (2018). High-density genotyping of the A.E. Watkins Collection of hexaploid landraces identifies a large molecular diversity compared to elite bread wheat. Plant Biotechnol J 16:165–175. 10.1111/pbi.12757.

Wingen, L.U., Orford, S., Goram, R., Leverington-Waite, M., Bilham, L., Patsiou, T.S., Ambrose, M., Dicks, J., and Griffiths, S. (2014). Establishing the A. E. Watkins landrace cultivar collection as a resource for systematic gene discovery in bread wheat. Theor Appl Genet 127:1831–1842. 10.1007/s00122-014-2344-5.

Zhang, B., Ma, L., Wu, B., Xing, Y., and Qiu, X. (2022). Introgression lines: valuable resources for functional genomics research and breeding in rice (*Oryza sativa* L.). Front Plant Sci 13:863789. 10.3389/fpls.2022.863789.

Zhao, X., Guo, Y., Kang, L., Yin, C., Bi, A., Xu, D., Zhang, Z., Zhang, J., Yang, X., Xu, J., et al. (2023). Population genomics unravels the Holocene history of bread wheat and its relatives. Nature plants 9:403–419. 10.1038/s41477-023-01367-3.

Zhu, T., Wang, L., Rimbert, H., Rodriguez, J.C., Deal, K.R., De Oliveira, R., Choulet, F., Keeble-Gagnere, G., Tibbits, J., Rogers, J., et al. (2021). Optical maps refine the bread wheat *Triticum aestivum* cv. Chinese Spring genome assembly. Plant J 107:303–314. 10.1111/tpj.15289.

Zimin, A.V., Marcais, G., Puiu, D., Roberts, M., Salzberg, S.L., and Yorke, J.A. (2013). The MaSuRCA genome assembler. Bioinformatics 29:2669–2677. 10.1093/bioinformatics/btt476.

