## Supplementary Figure for "Capturing the missing gene content of elite wheat by analysis of wheat landraces"

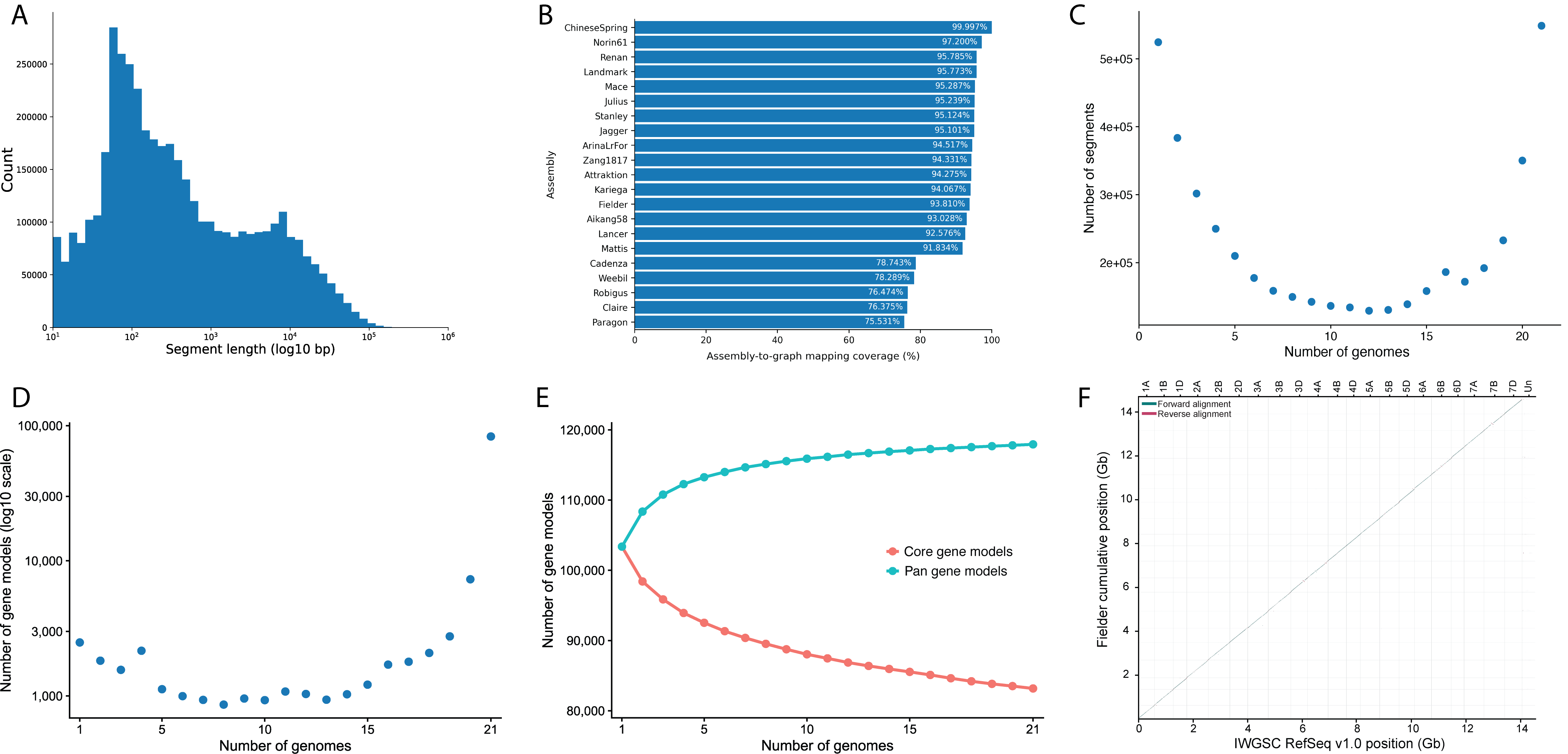

**Figure S1. Summary of the wheat graph pangenome constructed from 21 genome assemblies.** (**A**) Size distribution of all segments in the wheat graph pangenome (log scale). (**B**) Assembly-to-graph mapping coverage of the 21 wheat genome assemblies. (**C**) Number of segments shared by a number of individuals across 21 wheat genomes. (**D**) Number of predicted gene models shared by a number of individuals across 21 wheat genomes (log scale). (**E**) Pangenome and core-genome accumulation curves, with the upper and lower lines representing the total numbers of pangenome-predicted and core gene models, respectively. (F) Whole-genome alignment dot plot comparing the PacBio HiFi assembly of Fielder (y-axis) with the Chinese Spring RefSeq v1.0 (x-axis).

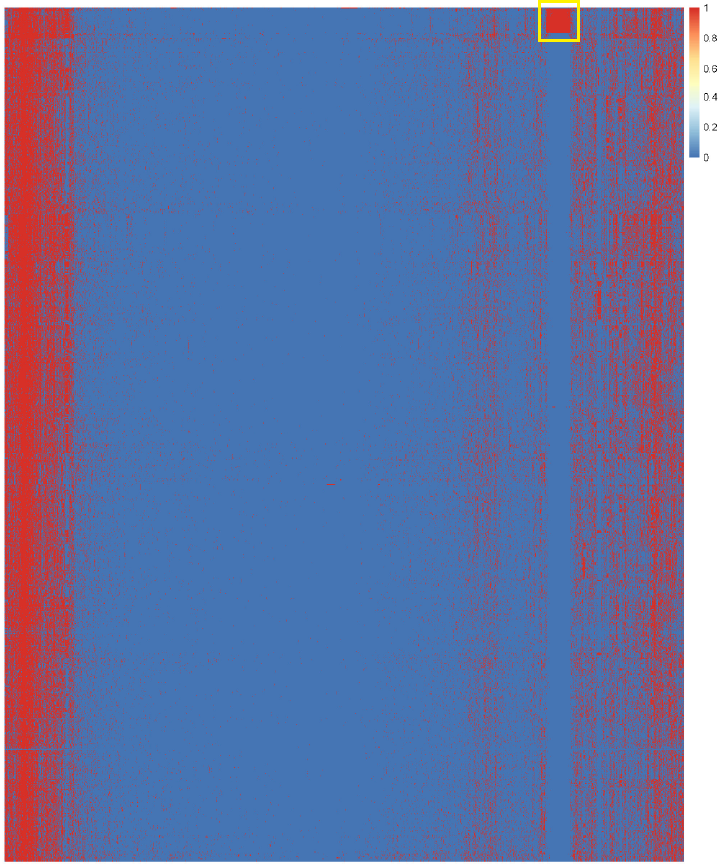

**Figure S2. Heat map of gene PAV of all gene models across all individuals.** Columns represent each individual and rows each gene model. Presence and absence of each gene model are represented by red and blue cells, respectively. The yellow box highlights the 581 gene models unique to 30 outlier individuals.

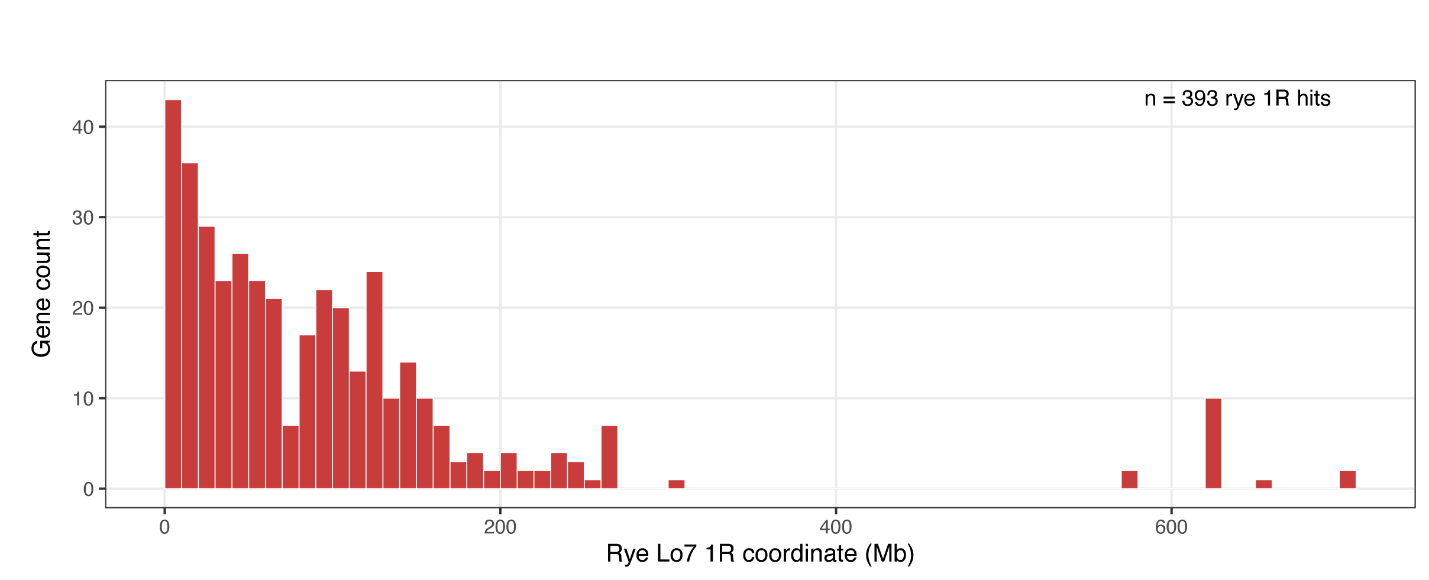

**Figure S3. Distribution of the genomic coordinates of 393 hits from the 581 outlier-specific gene models on rye Lo7 chromosome 1R.**

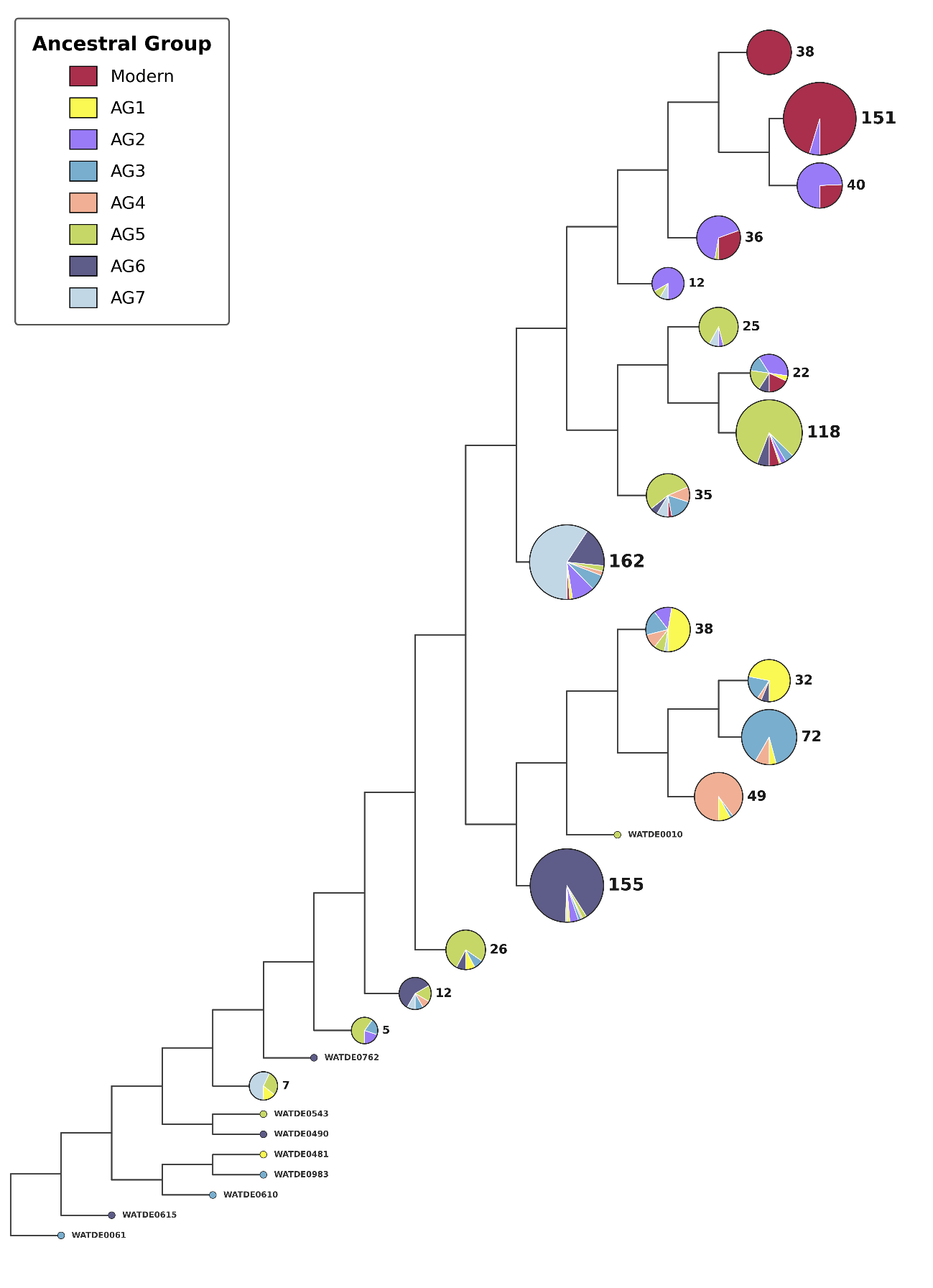

**Figure S4. Phylogenetic reconstruction based on gene PAV using Maximum Likelihood inference with 1,000 bootstrap replicates.** The tree is displayed as a cladogram, with pie charts at the tips showing the proportional composition of ancestral groups, coloured according to the same scheme as in Figure 1.

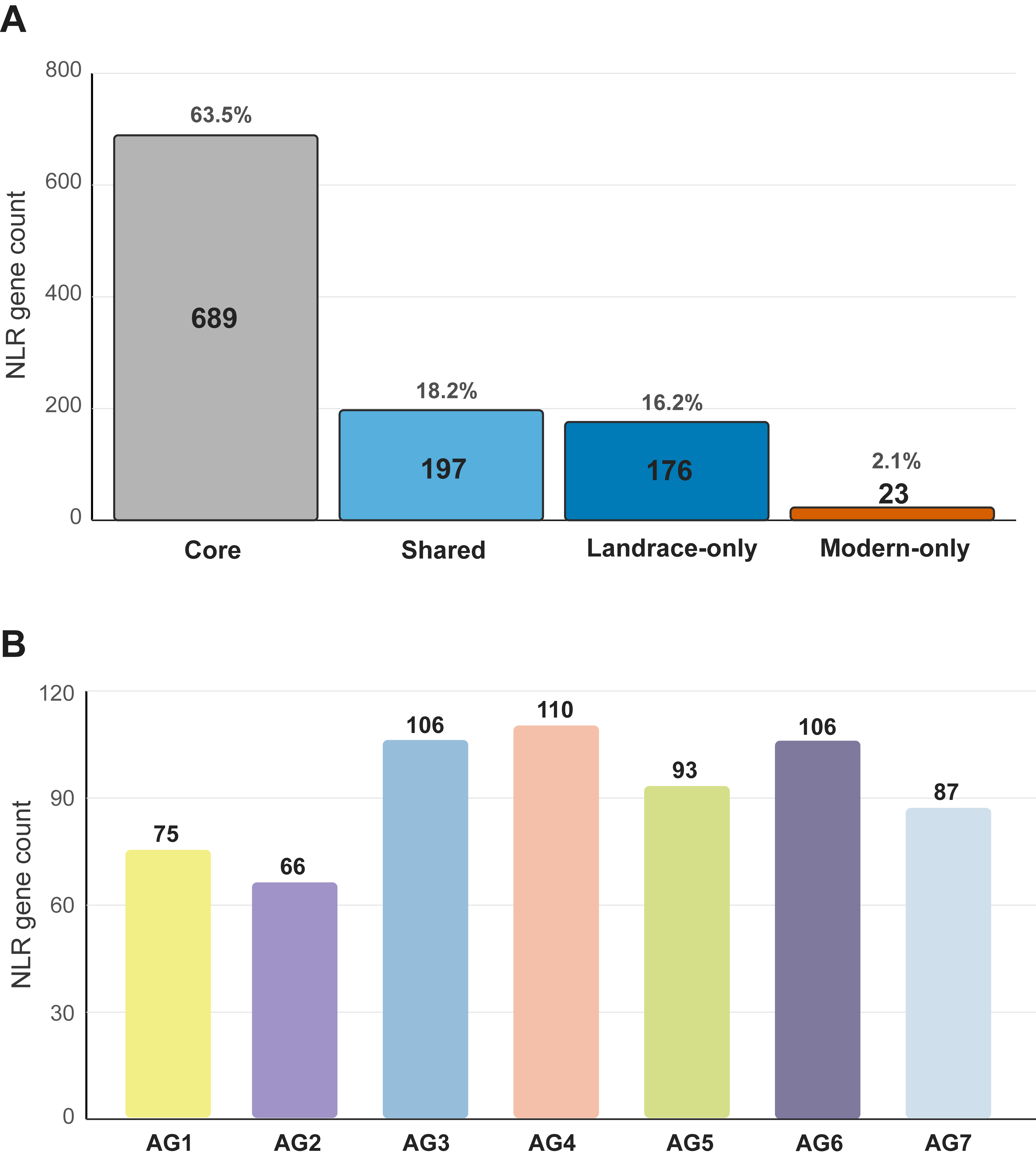

**Figure S5. NLR loci on the additional contigs and their presence across wheat landraces and modern cultivars.** (**A**) Presence categories of 1,085 NLR loci, including core (present in all seven landrace AGs and modern cultivars), shared (present in modern cultivars and at least one, but not all, landrace AGs), landrace-only, and Modern-only. (**B**) Distribution of landrace-specific NLR loci across the seven ancestral groups.
